# Developmental history modulates adult olfactory behavioral preferences via regulation of chemoreceptor expression in *C. elegans*

**DOI:** 10.1101/2022.06.07.495158

**Authors:** Travis Kyani-Rogers, Alison Philbrook, Ian G. McLachlan, Steven W. Flavell, Michael P. O’Donnell, Piali Sengupta

**Affiliations:** Department of Biology, Brandeis University, Waltham, MA 02454; Picower Institute for Learning and Memory, Department of Brain and Cognitive Sciences, Massachusetts Institute of Technology, Cambridge, MA 02139; Department of Molecular, Cellular and Developmental Biology, Yale University, New Haven, CT

## Abstract

Developmental experiences play critical roles in shaping adult physiology and behavior. We and others previously showed that adult *C. elegans* which transiently experienced dauer arrest during development (PD: post-dauer) exhibit distinct gene expression profiles as compared to control adults which bypassed the dauer stage. In particular, the expression patterns of subsets of chemoreceptor genes are markedly altered in PD adults. Whether altered chemoreceptor levels drive behavioral plasticity in PD adults is unknown. Here we show that PD adults exhibit enhanced attraction to a panel of food-related attractive volatile odorants including the bacterially-produced chemical diacetyl. Diacetyl-evoked responses in the AWA olfactory neuron pair are increased in both dauer larvae and PD adults, and we find that these increased responses are correlated with upregulation of the diacetyl receptor ODR-10 in AWA likely via both transcriptional and post-transcriptional mechanisms. We show that transcriptional upregulation of *odr-10* expression in dauer larvae is in part mediated by the DAF-16 FOXO transcription factor. Via transcriptional profiling of sorted populations of AWA neurons from control and PD adults, we further show that the expression of a subset of additional chemoreceptor genes in AWA is regulated similarly to *odr-10* in PD animals. Our results suggest that developmental experiences may be encoded at the level of olfactory receptor regulation, and provide a simple mechanism by which *C. elegans* is able to precisely modulate its behavioral preferences as a function of its current and past experiences.

## INTRODUCTION

Conditions experienced during early development have profound effects on adult phenotypes. Fetal malnutrition is a major risk factor for metabolic disorders in human adults, and adverse experiences in early life influence adult stress responses in many animal species (Alyamani and Murgatroyd, 2018; Cater and Majdic, 2021; de Gusmao Correia et al., 2012; Hanson and Gluckman, 2014; Smith and Ryckman, 2015). In addition to modulating general life history traits, early experiences can also affect specific adult behavioral phenotypes. For instance, exposure to an odorant during a critical developmental period has been shown to subsequently modulate the responses of adults to that odorant (Hong et al., 2017; Jin et al., 2016; Nevitt et al., 1994; Remy and Hobert, 2005). Adult phenotypic plasticity as a consequence of differential developmental experiences may allow adaptation to variable environments to optimize fitness (Sommer, 2020). Despite the prevalence and critical role of early experiences in shaping adult phenotypes, the underlying mechanisms are not fully understood.

*C. elegans* adults develop via one of two alternative developmental trajectories based on environmental conditions experienced during their first and second larval stages (Cassada and Russell, 1975). While larvae continue in the reproductive cycle through four larval stages (L1-L4) under favorable conditions, adverse conditions experienced during L1 instead drive larvae into the dauer diapause stage. Dauer larvae undergo extensive morphological, neuroanatomical and behavioral remodeling that maximizes their ability to survive harsh conditions, distinguishing them from their L3 larval counterparts (Albert and Riddle, 1983; Britz et al., 2021; Popham and Webster, 1979; Riddle, 1988). When growth conditions improve, they exit the dauer stage and resume development into reproductive adults. We and others previously showed that adult *C. elegans* that transiently passed through the dauer stage (henceforth referred to as post-dauer (PD) adults) differ markedly from adult animals that bypassed the dauer stage (referred to as control adults) in their life history traits including longevity, stress-resistance, and fecundity (Hall et al., 2010; Hall et al., 2013; Ow et al., 2018; Ow et al., 2021). This phenotypic plasticity is correlated with extensive changes in transcriptional profiles, as well as modification of the chromatin landscape (Bhattacharya et al., 2019; Hall et al., 2010; Hall et al., 2013; Ow et al., 2018; Vidal et al., 2018). Thus, isogenic populations of adult *C. elegans* hermaphrodites retain a molecular and phenotypic memory of their developmental history.

Food-seeking is a critical behavioral drive and is subject to extensive modulation as a function of an animal’s internal state and external conditions (Flavell et al., 2020; Heisler and Lam, 2017; Kim et al., 2017; Pool and Scott, 2014). Dauer larvae do not feed and the morphologies of their sensory neurons are extensively remodeled (Albert and Riddle, 1983; Britz et al., 2021; Popham and Webster, 1979; Riddle, 1988). Dauer larvae retain the ability to detect and respond to environmental cues including food-related chemical cues in order to assess whether conditions have improved sufficiently to trigger exit from the dauer stage and re-entry into the reproductive cycle. Chemosensory responses of *C. elegans* dauer larvae have been assessed in a limited set of studies and appear to be partly distinct from those of adults or L3 larvae (Albert and Riddle, 1983; Hallem et al., 2011; Vertiz et al., 2021; White et al., 2019). Whether PD adults retain these behavioral differences or exhibit further plasticity in food-seeking behaviors as a consequence of their distinctive developmental trajectory is largely unknown. The expression of many chemoreceptor genes in sensory neurons is altered in dauer larvae (Nolan et al., 2002; Peckol et al., 2001; Vidal et al., 2018), and the spatial patterns and/or levels of a subset of these genes is subsequently maintained or further altered in PD adults (Hall et al., 2010; Peckol et al., 2001; Vidal et al., 2018). Altered chemoreceptor expression profiles provide a plausible mechanism for chemosensory behavioral plasticity, but a correlation between changes in chemoreceptor gene expression and altered food-seeking behaviors in PD animals remains to be established.

Here we show that PD adults exhibit increased sensitivity to a panel of bacterially-produced attractive odorants including the chemical diacetyl, low concentrations of which are sensed by the AWA olfactory neuron pair. Consistently, both dauer larvae and PD adults also exhibit increased diacetyl-evoked responses in the AWA olfactory neurons. We find that the AWA-expressed *odr-10* diacetyl receptor gene is upregulated in dauer larvae; this upregulation is mediated in part via the DAF-16 FOXO transcription factor. While *odr-10* expression is subsequently downregulated in PD adults relative to levels in dauer larvae, levels of the ODR-10 receptor protein in the AWA cilium, the site of primary odorant transduction, are retained at higher levels in PD as compared to control adults. Via transcriptional profiling of sorted populations of AWA neurons from control and PD adults together with examination of expression from endogenous reporter-tagged alleles, we further find that the expression of a subset of additional AWA-expressed chemoreceptor genes is also upregulated in dauers, and is maintained at higher levels in PD adults. Together, our data demonstrate that altered chemoreceptor levels can underlie developmental stage- and history-dependent olfactory behavioral plasticity in *C. elegans*, and highlight the complexity of mechanisms regulating expression of individual chemoreceptor expression genes in this organism.

## MATERIALS and METHODS

### Strains and growth conditions

All *C. elegans* strains were maintained on nematode growth medium (NGM) seeded with *Escherichia coli* OP50 at 20°C unless stated otherwise. Dauer larvae were generated by picking 8-10 L4s from continuously growing animal populations onto 10 cm NGM plates and allowing growth and reproduction until food exhaustion (typically 7-9 days) at 25°C. Starved larvae were washed off plates with S-Basal buffer, and dauer larvae were selected by treatment with 1% SDS for 30 mins. To recover PD adults, a droplet containing 100-700 dauer larvae was pipetted onto a 10 cm NGM plate seeded with OP50 and allowed to grow for 48 hrs at 20°C. L4 larvae were picked onto fresh, seeded NGM plates 1 day prior to imaging.

For all experiments involving growth on plates containing auxin, NGM media was supplemented with the synthetic auxin 1mM 1-Naphthaleneacetic acid (NAA) from a 500 mM stock solution dissolved in 95% ethanol. Corresponding control NGM media was supplemented with an equal volume of 95% ethanol.

To collect adult animals that underwent L1 larval arrest, gravid adult hermaphrodites were bleached, and eggs were allowed to hatch overnight in M9 buffer containing 0.1% Triton X-100. 500-1000 L1 larvae were subsequently placed onto 10 cm NGM plates seeded with OP50 and grown at 20°C until adulthood.

### Genetics

All strains were constructed using standard genetic methods. Crosses were validated for the presence of the desired mutations using PCR-based amplification and/or sequencing. To generate strains for auxin-induced degradation specifically in AWA, animals were injected with a plasmid driving TIR1 under the AWA specific *gpa-4Δ6* promoter (PSAB1283: *gpa-4Δ6*p::*TIR1::SL2::mScarlet*; Table S1) at 5 ng/μl together with the *unc-122*p::*dsRed* co-injection marker at 50 ng/μl. A complete list of strains used in this work is available in Table S2.

### Molecular biology

Gene editing was performed using CRISPR/Cas9 and repair templates. Plasmids PSAB1279 (*odr-10::splitGFP_11_*) and PSAB1280 (*gpa-4Δ6*p::*splitGFP_1-10_*) (Table S1) were generated using traditional cloning from splitGFP constructs (KP#3315 and KP#3316, gift from J. Kaplan lab). An asymmetric donor template (Dokshin et al., 2018) was amplified from plasmid PSAB1279 to create long and short PCR fragments of 2308 and 600 bp, respectively, containing approximately 1 kb homology arms for CRISPR-mediated insertion. The asymmetric hybrid template was injected (each fragment at 250 ng/μl) together with crRNA (20 ng/μl; IDT Integrated DNA Technologies), tracrRNA (20 ng/μl: IDT), Cas9 protein (25 ng/μl; IDT), and *unc-122*p*::gfp* (40 ng/μl) as the co-injection marker. F1 animals expressing the injection marker were isolated. F2 progeny were screened by PCR for the insertion and confirmed by sequencing to obtain *odr-10(oy158)* (Table S2).

To mutate the predicted DAF-16 binding site upstream of *odr-10* in *odr-10(oy158)*, a conserved GTAAACA binding site 815 bp 5’ of the *odr-10* start codon (Wexler et al., 2020) was mutated to GTCCCCA to generate *odr-10(oy170)*. The injection mix contained an *odr-10* promoter repair template with the mutated site along with 32 bp 5’ and 3’ homology arms (590 ng/μl; IDT), *dpy-10* repair template (100 ng/μl, IDT), Cas9 protein (25 ng/μl; IDT), crRNA (20 ng/μl; IDT), and tracrRNA mix (20 ng/μl: IDT). F1 animals with *dpy* phenotypes were isolated and F1 and F2 progeny were screened by PCR and sequencing. Confirmed mutants were backcrossed to remove the *dpy-10* allele.

### Chemotaxis behavioral assays

Chemotaxis assays were performed essentially as described previously (Bargmann et al., 1993; Troemel et al., 1997). Behavioral attraction and avoidance assays were performed on 10 cm round or square plates, respectively. Each assay was performed in duplicate each day, and data are reported from biologically independent assays performed over at least 3 days. Behaviors of control and experimental animals were examined in parallel each day.

### Calcium imaging

Calcium imaging was performed essentially as previously described, using custom microfluidics devices (Chronis et al., 2007; Khan et al., 2022; Neal et al., 2015). Imaging was performed on an Olympus BX52WI microscope with a 40X oil objective and Hamamatsu Orca CCD camera. Video recordings were performed at 4 Hz. All odorants were diluted in filtered S-Basal buffer. 20 μM fluorescein was added to one buffer channel to confirm correct fluid flow in microfluidics devices. 1 mM (-)-tetramisole hydrochloride (Sigma L9756) was used to immobilize animals during imaging. To prevent animals from clogging the microfluidics loading arena and chip, 1 μl of poloxamer surfactant (Sigma P5556) was added to the S-Basal loading buffer. AWA neurons were imaged for one cycle of 30 sec buffer/30 sec odor/30 sec buffer, or for one cycle of 30 sec buffer/10 sec odor/20 sec buffer stimuli.

Recorded images were aligned with the template Matching plugin in Fiji (NIH) and cropped to include the AWA neuron soma and surrounding background fluorescence. The region of interest (ROI) was defined by outlining the AWA cell bodies, and an area of background fluorescence was chosen for background subtraction. To correct for photobleaching, an exponential decay was fit to the fluorescence intensity values for the first 30 sec and the last 20 sec of imaging. The resulting value was subtracted from original intensity values. Peak amplitude was calculated as the maximum change in fluorescence (F-F_0_) in the 10 sec following odor addition; F_0_ was set to the average ΔF/F_0_ value for 5 sec before odor onset. Data visualization was performed using RStudio (Version 1.4.1717). Photomask designs for customized adult and dauer microfluidic imaging chips were adapted from (Chronis et al., 2007) and are available at https://github.com/SenguptaLab/PDplasticity (also see below). AWA neuron mean baseline fluorescence (Figure S1C) was calculated by taking the average ΔF/F_0_ during the first 25 sec of imaging (0-25 sec) in each animal. Reported data were collected from biologically independent experiments over at least 2 days.

### Dauer microfluidics imaging device

Designs were based on olfactory imaging chips previously described (Chronis et al., 2007) with modifications to accommodate dauer larvae. To constrict the thinner dauer larvae in a similar manner to adults, it was necessary to reduce the cross-sectional area of the worm trap. This was accomplished by placing 10 μm-wide posts of decreasing width culminating in a 5 μm gap into which the worm nose was constricted (https://github.com/SenguptaLab/PDplasticity). Posts were used instead of fully narrowing the channel to preclude limiting fluid velocity while loading animals into the channel. AutoCAD drawings were used to generate an ink photomask (outputcity.com) which was subsequently used to generate master molds with 10 μm feature depth using negative photoresist SU-8 3005. To prevent delamination due to small features, a uniform 10 μm layer of SU-8 3005 photoresist was applied first to silicon wafers.

### Imaging and image analysis

Animals were mounted on 10% agarose pads set on microscope slides and immobilized using 10 mM (-)-tetramisole hydrochloride. Imaging for expression level quantification was performed on an inverted spinning disk confocal microscope (Zeiss Axiovert with a Yokogawa CSU22 spinning disk confocal head and a Photometerics Quantum SC 512 camera). Optical sections were acquired at 0.27 μM sections using either a 63X oil immersion objective when imaging neuron cell bodies, or a 100X oil immersion objective when imaging the cilia using SlideBook 6.0 software (Intelligent Imaging Innovations, 3i). *z*-projections of all optical sections at maximum intensity were generated using SlideBook 6.0 or FIJI/ImageJ (NIH).

Quantification of GFP or mNeonGreen levels in AWA soma was performed by tracing the ROI of each neuron cell body, and subtracting background fluorescence. The mean fluorescence of each cell body was calculated in FIJI. For quantification of ciliary ODR-10 levels in adult animals, an ROI of the cilia base and primary stalk was obtained and the integrated density of total fluorescence in the ROI was calculated using the corrected total cell fluorescence (CTCF) to take into account differences in the sizes of different ROIs and the highly branched structures of AWA cilia. Since the AWA cilia branches are collapsed in dauer larvae and are relatively smaller in L3 larvae, the ROI was traced around the entire cilia base and stalks. Three background ROIs were obtained and averaged to calculate the mean background fluorescence.

### Collection of AWA neurons

To obtain control animals, adult hermaphrodites expressing the stably integrated transgene *gpa-4Δ6*p*::myrGFP* (PY10421, Table S2) were bleached, and eggs allowed to hatch in M9 with 0.1% Triton-100. 50,000-100,000 growth-synchronized L1 larvae were plated onto 15 cm 8P growth plates seeded with 1 ml of an overnight culture of *E. coli* NA22 grown in 2XYT media. L1 larvae were allowed to grow for 40-48 hrs at 20°C to obtain populations of L4 larvae. To obtain PD animals, SDS-selected dauer larvae were plated onto 15 cm 8P plates seeded with 1 ml of an overnight culture of *E. coli* NA22 grown in 2XYT media, and allowed to recover to the L4 stage for 20-24 hrs at 20°C.

Animals were collected and cells dissociated essentially as described previously (Taylor et al., 2021). In brief, animals were washed off plates with M9 buffer and incubated in lysis buffer (200 mM Dithiothreitol (DTT), 0.25% Sodium dodecyl sulfate (SDS), 20 mM HEPES buffer (4-(2-hydroxyethyl)-1-piperazineethanesulfonic acid), 3% sucrose) for 5 mins. Lysed animals were washed 5X with egg buffer (118 mM NaCl, 48 mM KCl, 2 mM CaCl_2_, 2 mM MgCl_2_, 25 mM HEPES, pH 7.3) following which the worm pellet was resuspended in 750 μl of 15 mg/ml Pronase (Sigma P8811) in egg buffer. To promote dissociation, the mixture was pipetted frequently for 20 mins and progress of dissociation was monitored under a light microscope. Pronase digestion was terminated by adding L-15-10 media (L-15 media + 10% volume FBS, HI) at 4°C. Dissociated worms were centrifuged at 4°C and the worm pellet was resuspended in egg buffer and recentrifuged at 4°C. The supernatant was filtered through a 35 µm filter in preparation for FACS sorting. A 25 μl aliquot of the dissociated worm pellet was pipetted directly into Trizol for analysis as the whole worm sample.

Prior to performing FACS, 0.5 μl of 1 mg/ml DAPI was added to 1 ml of the cell suspension to exclude dead cells. AWA neurons were captured by isolating the high GFP+ and low DAPI compartment and cells were sorted directly into Trizol. Cells from this compartment were also sorted onto glass slides and imaged under a fluorescence light microscope to confirm the presence of intact GFP+ cells. Following cell collection, collection tubes were briefly spun down and frozen at -80°C. 1,000 - 10,000 cells were isolated per sorting run.

### RNA sequencing

RNA was extracted from collection tubes using standard chloroform and isopropanol precipitation. The RNA pellet was resuspended in RNAase-free water and any DNA digested using the RNase-Free DNase kit (Qiagen). RNA was eluted using the RNeasy MinElute Cleanup Kit (Qiagen). mRNA was isolated and libraries amplified using Smart-Seq v4 and library quality checked with Bioanalyzer using Agilent High Sensitivity Gel. Libraries were sequenced on Illumina NextSeq500. Library preparation and sequencing was performed at the MIT BioMicro Center (https://biology.mit.edu/tile/biomicro-center/).

Raw RNA-Seq reads were quality checked, adapter trimmed, and aligned to the *C. elegans* genome using STAR ALIGNER (https://github.com/alexdobin/STAR). Principal component analyses were performed on the normalized read counts of the 10,000 most variable genes across all samples. Differential expression analysis was performed using DESeq2 with lfc shrinkage on. Differential expression was determined with a significance cut-off of 0.05 (adjusted *p*-value; Wald test with Benjamini-Hochberg corrections) and the indicated Log_2_ fold change cut off of either -1.5 and 1.5 or -2 and 2. AWA enrichment analysis was performed using the web-based Tissue Enrichment Analysis tool with default parameters (https://www.wormbase.org/tools/enrichment/tea/tea.cgi) (Angeles-Albores et al., 2016).

### Statistical analyses

Normal distribution of the data was assessed using the Shapiro-Wilks test (GraphPad Prism v9.0.2). Parametric data were analyzed using a two-tailed Welch’s t-test or ANOVA. Non-parametric data were analyzed using a Mann-Whitney t-test or Kruskal-Wallis test across multiple conditions. Post hoc corrections for multiple comparisons were applied to data in which more than two groups were analyzed. The specific tests used and corrections applied are indicated in each Figure Legend.

## RESULTS

### Post-dauer adult animals exhibit enhanced attraction to a subset of volatile odorants

*C. elegans* adult animals are strongly attracted to a subset of chemicals that indicates the presence of nutritious bacteria, the major food source for these nematodes (Ferkey et al., 2021). Attractive volatile odorants are primarily sensed by the AWA and AWC olfactory neurons pairs in the bilateral head amphid organs of *C. elegans* (Bargmann et al., 1993). Consistently, adult hermaphrodite animals grown continuously under favorable environmental conditions (control adults; Figure 1A) exhibited robust attraction to a range of concentrations of odorants sensed by the AWA and/or AWC neurons (Figure 1B).

**Figure 1.**
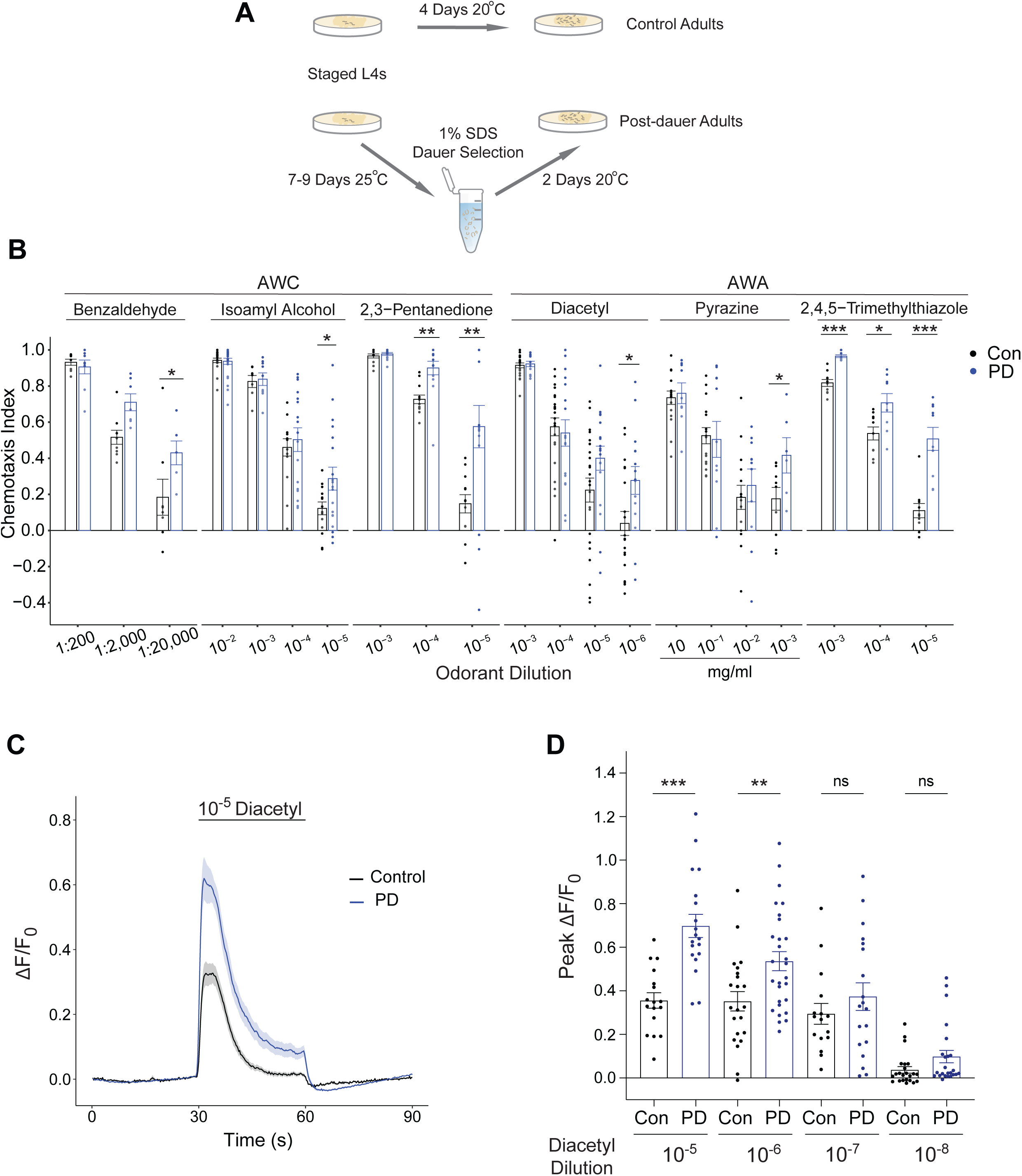
Olfactory responses to a panel of volatile attractants is increased in PD adults. **A)** Cartoon of growth conditions for the generation of control and post-dauer (PD) adults. **B)** Behavioral responses of wild-type control and PD adults to a panel of volatile odorants at the indicated dilutions. Chemotaxis index = (Number of animals at the odorant – number of animals at the diluent ethanol) / (Number of animals at the odorant + number of animals at ethanol). Each dot is the chemotaxis index of a single assay plate containing ∼50-300 adult hermaphrodites. Bars represent the mean; error bars are SEM. The behaviors of control and PD animals were assayed in parallel in duplicate; ≥3 independent experiments. *, **, *** indicate different at *P*<0.05, 0.01, and 0.001, respectively (two-tailed Welch’s t-test). **C)** Average changes in GCaMP2.2b fluorescence in AWA soma to a 30 sec pulse of 10^-5^ diacetyl in control and PD adults. Shaded regions indicate SEM. n ≥ 16 animals (1 neuron per animal) each. **D)** Quantification of peak fluorescence intensity changes in AWA soma expressing GCaMP2.2b to a 30 sec pulse of diacetyl at the indicated concentrations. Bars represent mean, error bars are SEM. n ≥ 16 animals (1 neuron per animal) each. Control and PD adults were examined in parallel over at least three days. ** and *** indicate different at each concentration at *P*<0.01 and 0.001, respectively (two-tailed Welch’s t-test); ns – not significant.

To test whether passage through the dauer stage influences the olfactory preference behaviors of adult animals, we grew wild-type animals under food-restricted conditions to promote entry into the dauer stage as previously described (Figure 1A) (Ow et al., 2018). Dauer larvae were collected via SDS-mediated selection and subsequently allowed to resume reproductive growth on bacterial food (Figure 1A). Similar to control adults, young post-dauer (henceforth referred to as PD) adult animals were also robustly attracted to both AWA- and AWC-sensed volatile odorants (Figure 1B). We found that PD adults typically exhibited enhanced attraction to lower odorant concentrations relative to control adults (Figure 1B). Animals that experienced starvation-induced developmental arrest at the L1 larval stage did not exhibit similar enhanced attraction responses as adults (Figure S1A). PD animals have previously been shown to exhibit decreased avoidance of the aversive pheromone ascr#3 (Sims et al., 2016). Control and PD animals avoided high concentrations of benzaldehyde to a similar extent (Figure S1B) although the aversion responses of PD adults to high concentrations of octanol were weakly decreased (Figure S1B). We conclude that the behavioral responses of adult animals to multiple attractive volatile chemicals are sensitized upon passage through the dauer diapause stage.

We next tested whether increased behavioral attraction correlates with enhanced odorant responses in olfactory neurons by examining odorant-evoked changes in intraneuronal calcium dynamics using a genetically encoded calcium indicator. We focused our attention on the odorant diacetyl, low concentrations of which are sensed by the AWA olfactory neuron pair (Bargmann et al., 1993), and for which the cognate receptor and signal transduction mechanisms have been described (see below) (Colbert et al., 1997; Roayaie et al., 1998; Sengupta et al., 1996). Low concentrations of diacetyl consistently evoked responses of larger amplitude in the AWA neurons of PD as compared to control adults consistent with the observed increased behavioral responses to this chemical (Figure 1C-D). This amplitude difference is unlikely to arise simply due to differences in the expression levels of the AWA-expressed calcium sensor (Figure S1C). Together, these data indicate that neuronal responses of adult animals to the attractant diacetyl are enhanced upon passage through the dauer stage.

### Ciliary levels of the diacetyl receptor ODR-10 are increased as a function of developmental history

We and others previously reported that control and PD animals exhibit significant differences in gene expression profiles including in individual sensory neurons (Bhattacharya et al., 2019; Hall et al., 2010; Sims et al., 2016; Vidal et al., 2018). Each chemosensory neuron type in *C. elegans* expresses multiple GPCRs, expression of a subset of which has previously been shown to be regulated by external and internal state (Gruner et al., 2014; Lanjuin and Sengupta, 2002; Nolan et al., 2002; Peckol et al., 2001; Ryan et al., 2014; Troemel et al., 1995; Vidal et al., 2018). Low concentrations of diacetyl are sensed by the ODR-10 olfactory receptor which is expressed specifically in the AWA neurons and localizes to their sensory cilia, the primary site of olfactory signal transduction (Sengupta et al., 1996). Behavioral and neuronal responses to diacetyl have previously been shown to be regulated by feeding state, developmental stage and somatic sex via expression changes in *odr-10* (Ryan et al., 2014; Wexler et al., 2020), but whether *odr-10* expression is also modulated via dauer passage is unclear. Since state-dependent regulation of individual olfactory receptor genes provides a simple mechanism for mediating odorant-selective behavioral plasticity, we tested the hypothesis that altered expression of *odr-10* underlies the observed plasticity in diacetyl responses in PD adults.

An endogenous *odr-10* allele tagged with *t2A::mNeonGreen* (*odr-10::t2A::mNG*) was specifically expressed in both AWA neurons (Figure 2A) (McLachlan et al., 2022). However, expression levels of the fluorescent reporter protein were not significantly altered in PD as compared to control adults (Figure 2B). Since this reporter does not allow assessment of ciliary ODR-10 protein levels, we next quantified ODR-10 protein levels in AWA cilia from the endogenous *odr-10* allele tagged with the split-GFP reporter GFP_11_ (*odr-10(oy158))*, and reconstitution of GFP via expression of the GFP_1-10_ fragment under a constitutive AWA-specific promoter (Kamiyama et al., 2016). We confirmed that tagging *odr-10* with GFP_11_ had no effect on behavioral responses to diacetyl, and that these animals continued to exhibit enhanced attraction to diacetyl upon passage through the dauer stage (Figure S2A). The reconstituted ODR-10 fusion protein was localized to the extensively branched AWA cilia in both control and PD animals (Figure 2C). Since the complex architecture of the AWA cilium precluded precise quantification of overall ciliary protein levels in adult animals, we restricted our analysis to assessing ODR-10 protein levels at the ciliary base and in the primary ciliary stalk. We found that reconstituted ODR-10::GFP levels in AWA cilia were significantly higher in PD than in control animals (Figure 2C-D). These observations raise the possibility that increased ciliary levels of ODR-10 protein in PD animals may contribute to the increased diacetyl responses of PD animals.

**Figure 2.**
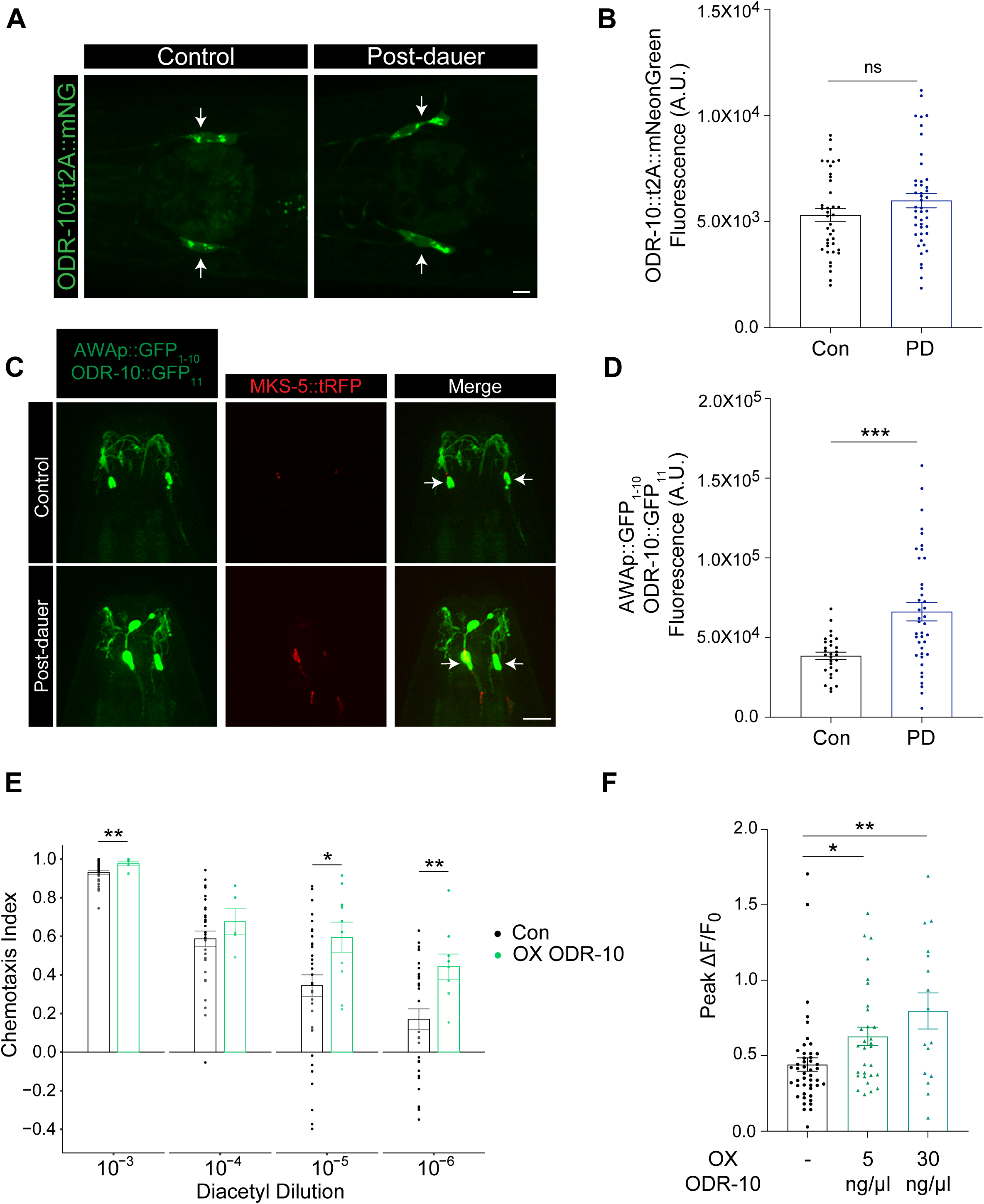
Ciliary ODR-10 levels are increased in PD adults. **A)** Representative images of AWA neurons expressing *odr-10*::*t2A*::*mNeonGreen* from the endogenous *odr-10* locus in control and PD adults. Arrows indicate AWA cell bodies. Anterior is at left. Scale bars: 5 µm. **B)** Quantification of ODR-10::t2A::mNeonGreen fluorescence in AWA soma in control and PD adults. Each dot is a measurement from an individual neuron. Bars represent the mean; error bars are SEM. n ≥ 39 neurons (≥ 21 animals). Control and PD adults were imaged in parallel on the same day; ≥3 independent experiments. ns – not significant. **C)** Representative images of ODR-10::splitGFP_11_ expression in AWA cilia of control and PD adults also expressing *gpa-4Δ6*p::splitGFP_1-10_. MKS-5::tagRFP marks the ciliary transition zones. Arrows indicate the ciliary base. Anterior is at top. Scale bars: 5 µm. **D)** Quantification of total ciliary ODR-10::splitGFP_11_ fluorescence in AWA cilia base and primary stalk in control and PD adults also expressing *gpa-4Δ6*p::splitGFP_1-10_. Each dot is a measurement from an individual animal. Bars represent the mean, error bars are SEM. n ≥ 29 animals. Control and PD adults were imaged in parallel; ≥3 independent experiments. *** indicates different at *P*<0.001 (two-tailed Welch’s t-test). **E)** Behavioral responses of wild-type control and wild-type control animals overexpressing ODR-10::tagRFP under the *gpa-4Δ6* promoter (OX ODR-10; 5 ng/ml) to the indicated dilutions of diacetyl. Each dot represents the chemotaxis index of a single assay plate containing ∼50-300 adult hermaphrodites. Bars represent the mean; error bars are SEM. A subset of control and experimental chemotaxis assays were performed in parallel over at least three days. * and ** indicate different at each concentration at *P*<0.05 and <0.01, respectively (two-tailed Welch’s t-test). **F)** Quantification of peak fluorescence intensity changes in AWA to a 10 sec pulse of 10^-7^ diacetyl. Responses of wild-type control animals injected with *gpa-4Δ6*p*::odr-10::tagRFP* at 5 and 30 ng/ml are shown. Bars represent mean, error bars are SEM. n ≥ 16 animals (1 neuron per animal) each. Animals were examined over two days with the exception of OX ODR-10 (30 ng/μl). * and ** indicate different at *P*<0.05 and <0.01, respectively (Kruskal-Wallis with Dunn’s multiple comparisons test).

To establish whether increasing ODR-10 levels alone in AWA is sufficient to enhance diacetyl response sensitivity even in control animals, we overexpressed *odr-10* from the constitutive *gpa-4Δ6* promoter and examined diacetyl responses via both behavioral assays and imaging of diacetyl-evoked intracellular calcium dynamics. As shown in Figure 2E, the behavioral responses of control animals overexpressing *odr-10* were higher than those of wild-type control animals across multiple concentrations of diacetyl. Overexpression of *odr-10* was also sufficient to increase diacetyl-evoked calcium responses in AWA in control adults (Figure 2F, Figure S2B). We conclude that increased ODR-10 levels may be sufficient to account for the enhanced diacetyl responses of PD animals.

### *odr-10* expression and diacetyl behavioral responses are increased in dauer larvae

*C. elegans* dauer larvae as well as the analogous infective juvenile larvae of parasitic nematodes have been shown to exhibit distinct olfactory behaviors (Vertiz et al., 2021). The expression of many neuronal genes including chemoreceptor genes is also markedly altered in dauer larvae (Bhattacharya et al., 2019; Hall et al., 2010; Nolan et al., 2002; Peckol et al., 2001; Vidal et al., 2018). The dauer-specific expression patterns of a subset of these genes is maintained in PD adults, while the expression of other genes is further altered to a PD-specific pattern or restored to the pattern observed in control adults (Hall et al., 2010; Peckol et al., 2001; Vidal et al., 2018). We asked whether *odr-10* expression is modulated in dauers, following which ciliary protein levels may be subsequently maintained at higher levels in PD animals.

Expression of the *odr-10::t2A::mNG* reporter remained restricted to the AWA neurons in dauer larvae (Figure 3A). In contrast to our observations in PD adults, *odr-10::t2A::mNG* expression was strongly upregulated in dauer animals as compared to levels in L3 larvae (Figure 3A-B). Consistently, reconstituted ODR-10::GFP protein levels were also increased in AWA cilia in dauers (Figure 3C-D).

**Figure 3.**
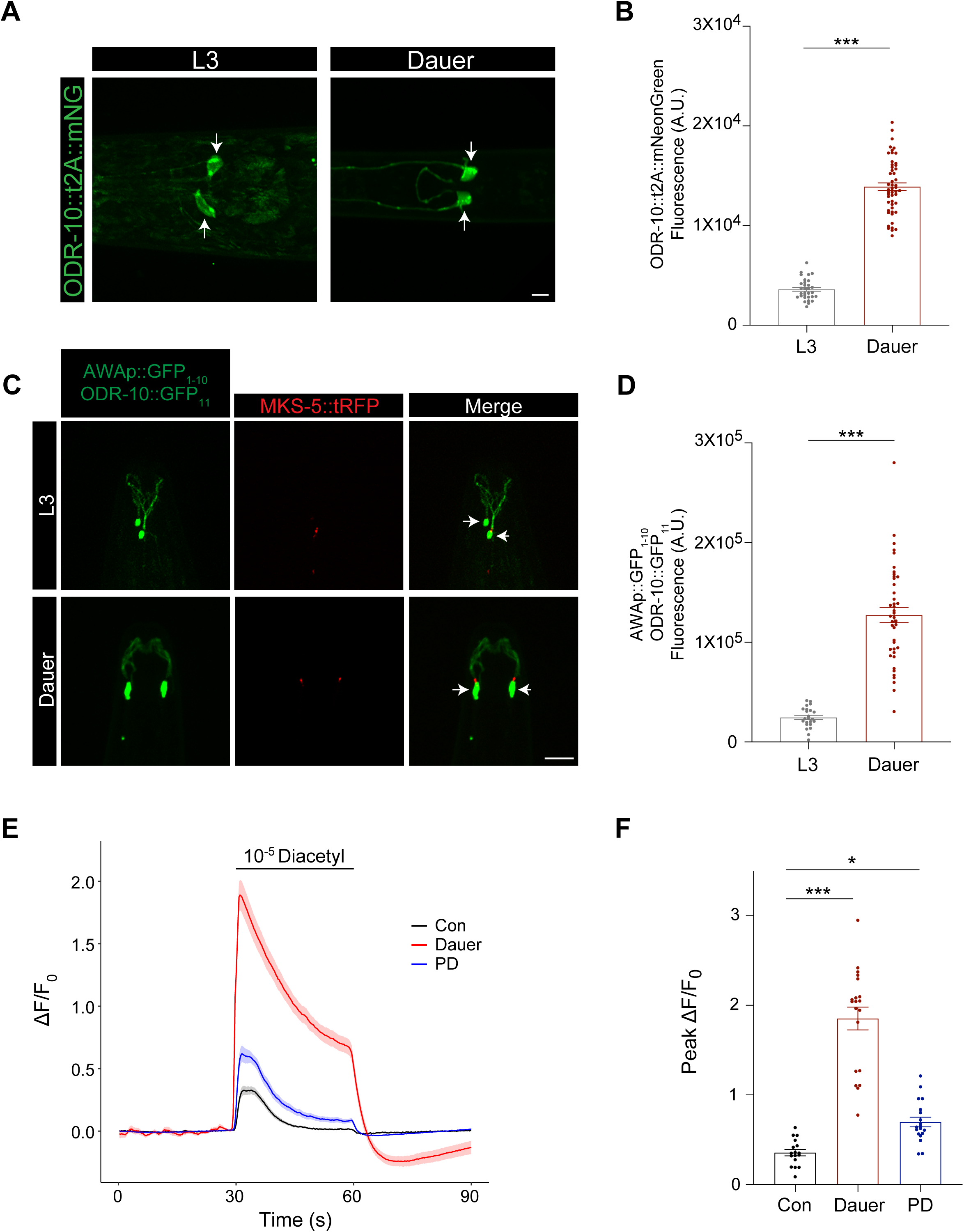
Dauer larvae exhibit upregulated *odr-10* expression and enhanced diacetyl responses. **A)** Representative images of AWA neurons in L3 and dauer larvae expressing ODR-10::t2A::mNeonGreen from the endogenous *odr-10* locus. Arrows indicate AWA cell bodies. Anterior is at left. Scale bar: 5 µm. **B)** Quantification of ODR-10::t2A::mNeonGreen fluorescence in AWA neurons of the indicated animals. Each dot is a measurement from a single neuron. Bars represent the mean; error bars are SEM. n ≥ 31 neurons; (≥ 18 animals). L3 and dauer larvae were imaged on the same day, ≥3 independent experiments. *** indicates different between indicated at *P*<0.001 (two-tailed Welch’s t-test). **C)** Representative images of reconstituted ODR-10::GFP expression in AWA cilia of L3 and dauer larvae. MKS-5::tagRFP marks the ciliary transition zones. Arrows indicate the AWA cilia base. Anterior at top. Scale bars: 5 µm. **D)** Quantification of total reconstituted ODR-10::GFP fluorescence in the AWA ciliary base and primary stalk in L3 and dauer animals. Each dot is a measurement from an individual animal. Bars represent the mean total fluorescence, error bars are SEM. n ≥ 23 animals. *** indicates different between indicated at *P*<0.001 (two-tailed Welch’s t-test). **E)** Average changes in GCaMP2.2b fluorescence in AWA to a 30 sec pulse of 10^-5^ dilution of diacetyl in control and PD adults, and dauer larvae. Shaded regions indicate SEM. Control and PD adult data are repeated from Figure 1D. n ≥ 16 neurons (1 neuron per animal). **F)** Quantification of peak fluorescence intensity changes in AWA expressing GCaMP2.2b to a 30 sec pulse of 10^-5^ dilution of diacetyl. Each dot is a measurement from a single neuron. Bars represent mean fluorescence, error bars are SEM. n ≥ 16 neurons (1 neuron per animal). Control and PD adult data are repeated from Figure 1D. * and *** indicate different at *P*<0.05 and <0.001, respectively (one-way ANOVA with Tukey’s multiple comparisons test).

We tested whether upregulation of *odr-10* expression in dauers correlates with increased diacetyl responses in these animals. To examine diacetyl-evoked calcium responses in AWA, we modified the microfluidics imaging device typically used for imaging *C. elegans* adults (Chronis et al., 2007) to accommodate the thinner and smaller dauer larvae (see Materials and Methods), although we were unable to use these or the adult imaging devices to examine L3 larvae. In response to a pulse of 10^-5^ dilution of diacetyl, dauer animals exhibited markedly increased responses as compared to control or PD adults (Figure 3E-F). Altered expression levels of the calcium sensor in AWA are unlikely to account for the observed increase in diacetyl responses in dauer larvae (Figure S1C). We could not reliably assess the behavioral responses of dauers to diacetyl in our behavioral assays in part due to their unique locomotory patterns (Bhattacharya et al., 2019; Gaglia and Kenyon, 2009), although we note that a previous report indicated that dauers exhibit decreased behavioral responses to diacetyl (Vertiz et al., 2021). We conclude that *odr-10* may be upregulated upon entry into the dauer stage possibly via transcriptional mechanisms and correlates with increased diacetyl responses. While this transcriptional upregulation is not maintained in PD adults, increased ODR-10 protein levels in AWA cilia correlates with enhanced diacetyl responses in PD adults.

### Upregulation of *odr-10* expression in dauer larvae is mediated in part via the DAF-16 FOXO transcription factor

The TGF-β and insulin signaling pathways act in parallel to regulate dauer formation in response to adverse environmental conditions (Fielenbach and Antebi, 2008; Riddle and Albert, 1997). Insulin signaling inhibits nuclear translocation of the DAF-16 FOXO transcription factor, and DAF-16 is nuclear-localized in dauer larvae (Aghayeva et al., 2021; Lin et al., 1997; Ogg et al., 1997). Since this molecule has been implicated in the altered regulation of sensory gene expression in dauer larvae (Aghayeva et al., 2021; Bhattacharya et al., 2019; Wexler et al., 2020), we tested whether the observed upregulation of *odr-10* expression in dauers is mediated in part via DAF-16-dependent transcriptional regulation.

Since *daf-16* mutants cannot enter the dauer stage, we assessed the effects of *daf-16* depletion via auxin-induced degradation specifically in AWA (Nishimura et al., 2009; Zhang et al., 2015). We expressed the auxin receptor TIR1 in AWA in a strain in which the endogenous *daf-16* locus has been edited to include a degron tag (Aghayeva et al., 2020; Aghayeva et al., 2021). To obtain larger numbers of dauer larvae, these experiments were performed in *daf-2* insulin receptor mutants that constitutively enter the dauer stage even under favorable environmental conditions (Figure 4A) (Gems et al., 1998; Riddle et al., 1981). As in wild-type animals, *odr-10::t2A::mNG* expression levels were upregulated in *daf-2* dauer larvae as compared to expression levels in control or PD *daf-2* adults in the absence of auxin treatment (Figure 4B). We found that auxin-mediated depletion of DAF-16 in AWA decreased, although did not fully abolish, the upregulated *odr-10* expression observed in dauer animals (Figure 4Ai, 4B). Growth on auxin had little effect on *odr-10* expression levels in adult animals that bypassed the dauer stage (Figure 4Aii, 4B), although we note that *odr-10* was previously identified as a putative DAF-16-regulated gene in a comparison of the neuronal transcriptomes of *daf-2* and *daf-16; daf-2* adult animals (Kaletsky et al., 2015). Addition of auxin only during dauer recovery also did not affect *odr-10* expression in adult animals (Figure 4Aiii, 4B). We infer that DAF-function in AWA is partly necessary during dauer entry to transcriptionally upregulate *odr-10* expression, although we are unable to exclude the possibility that DAF-16 is not fully depleted in AWA under these conditions.

**Figure 4.**
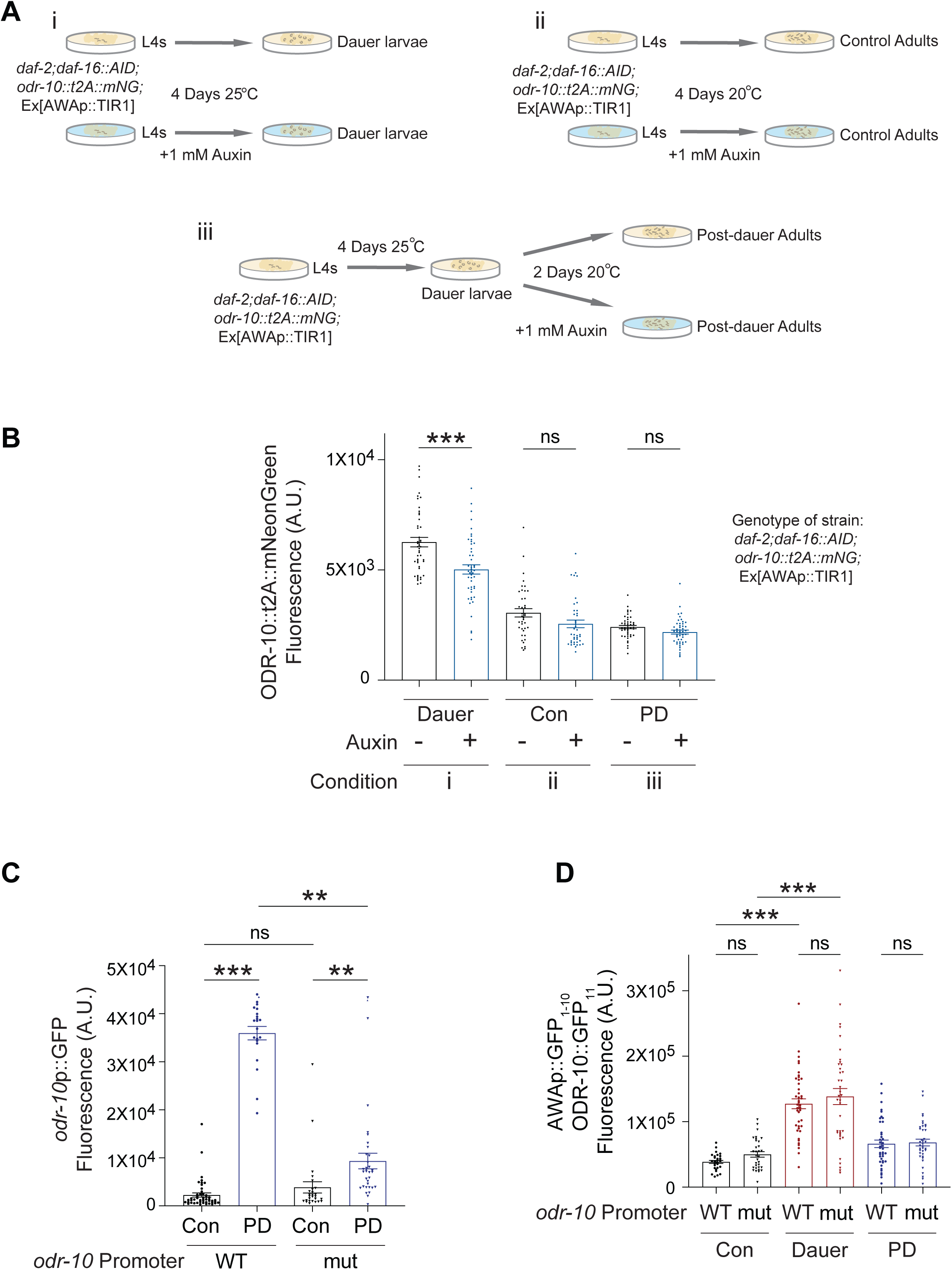
DAF-16 FOXO may be partly necessary for upregulation of *odr-10* expression in dauer larvae. **A)** Cartoons of different growth conditions of *daf-2(e1368)* animals expressing *daf-16::mNeptune::AID, odr-10::t2A::mNeonGreen* and *gpa-4Δ6*p::TIR1 with or without 1 mM auxin. **B)** Quantification of mean *odr-10::t2A::mNeonGreen* fluorescence in AWA in animals grown with or without 1 mM auxin in conditions indicated in **Ai-Aiii**. Each point is a measurement from an individual AWA neuron. Bars represent mean fluorescence, error bars are SEM. n ≥ 46 neurons (≥ 23 animals) for dauers; n ≥ 40 neurons (≥ 22 animals) for adults. Dauer larvae and control and PD adults grown with or without 1 mM auxin were imaged in parallel; 3 independent experiments. *** indicates different between indicated at *P*<0.001 (two-tailed Welch’s t-test); ns – not significant. **C)** Quantification of GFP fluorescence in AWA neurons expressing GFP under 1.0 kb of wild-type or mutant *odr-10* promoter sequences in which a conserved DAF-16 binding site has been mutated (mut) (Wexler et al., 2020). Each point is a measurement from an individual neuron. Bars represent mean fluorescence, error bars are SEM. n ≥ 21 neurons (≥ 27 animals). ** and *** indicate different at *P*<0.01 and <0.001, respectively (Kruskal-Wallis test with Dunn’s multiple comparisons test); ns – not significant. **D)** Quantification of total reconstituted ODR-10::GFP fluorescence in AWA ciliary base and stalk. mut indicates animals in which a conserved DAF-16 binding sequence within the endogenous *odr-10* promoter has been mutated. Each dot is a measurement from an individual neuron. Bars represent the mean total fluorescence, error bars are SEM. n ≥ 29 animals. Control and PD wild-type data are repeated from Figure 2D. *** indicates different at *P*<0.001 (one-way ANOVA with Tukey’s multiple comparison test); ns – not significant.

A putative DAF-16 binding site in the proximal regulatory sequences of *odr-10* was previously shown to be necessary for starvation-dependent upregulation of *odr-10* expression driven from a transcriptional reporter in *C. elegans* males (Wexler et al., 2020). In contrast to the expression of the endogenous *odr-10::t2A::mNG* reporter, transgenic expression of GFP driven by ∼1kb of the *odr-10* promoter containing the predicted DAF-16 binding site retained high levels of expression in PD adults (Figure 4C). Mutating the predicted DAF-16 binding site in this reporter construct decreased expression levels in PD adults (Figure 4C). However, mutating this site in the endogenous *odr-10* locus via gene editing had no effect on the upregulated levels of ciliary ODR-10 protein in either dauer or PD animals (Figure 4D). These results suggest that multiple DAF-16 binding sites in *odr-10* regulatory sequences may contribute redundantly in the context of the endogenous promoter to the upregulation of *odr-10* expression in dauer larvae. Alternatively, DAF-16 may act indirectly to regulate endogenous *odr-10* expression.

### AWA neurons exhibit distinct gene expression profiles in control and PD adults

In addition to diacetyl, PD adults also exhibit increased responses to the AWA-sensed odorants pyrazine and 2,4,5-trimethylthiazole (Figure 1B). Although the receptors for these chemicals are as yet unidentified, this observation suggests that the expression of chemoreceptors in addition to *odr-10* in AWA may also be altered as a function of dauer passage. To test this notion, we dissociated control and PD L4 larvae expressing GFP specifically in AWA, collected GFP-labeled populations of AWA neurons via fluorescence-activated cell sorting (FACS), and performed transcriptional profiling (Taylor et al., 2021). In order to obtain large populations of growth-synchronized L4 animals for cell sorting, the control population was grown from L1-arrested larvae (see Materials and Methods). In parallel, we also transcriptionally profiled populations of dissociated but unsorted cells from control and PD L4 animals.

Principal component analyses indicated that with the exception of one sample, the RNA-Seq profiles of all biologically independent replicates of sorted control and PD AWA neurons were present in a cluster distinct from that obtained from cells collected from whole animals (Figure S3A). AWA-expressed genes were the most enriched in the dataset collected from sorted AWA neurons (Figure S3B). Moreover, the 20 most highly expressed genes in AWA as predicted by the CeNGEN neuronal profiling project (Taylor et al., 2021) were robustly represented as upregulated in the AWA RNA-Seq data as compared to data from whole animals (Figure S3C). These observations indicate that we successfully enriched and profiled populations of AWA neurons.

Comparison of the gene expression changes in the datasets from control and PD whole animals indicated that the expression of multiple G protein-coupled receptor (GPCR) genes was altered as a function of dauer passage (Figure 5A-B, File S1). There was minimal overlap with previously published datasets of genes differentially regulated in PD vs control whole animals (Hall et al., 2010; Ow et al., 2018), likely due to differences in conditions of animal growth and sample preparation although we note that genes predicted to be involved in GPCR signaling were also previously identified as a differentially expressed category (Hall et al., 2010). Up- or downregulated chemoreceptor genes are predicted to be expressed in multiple chemosensory neuron types, indicating that modulation of chemoreceptor gene expression as a function of developmental history is mediated at the level of individual chemoreceptor genes and not sensory neuron types. However, despite the established importance of neuropeptide and hormonal signaling in regulating dauer entry (Fielenbach and Antebi, 2008; Riddle and Albert, 1997), the expression of only a few predicted neuropeptide genes appeared to be altered in PD as compared to control animals (Figure 5B, File S1).

**Figure 5.**
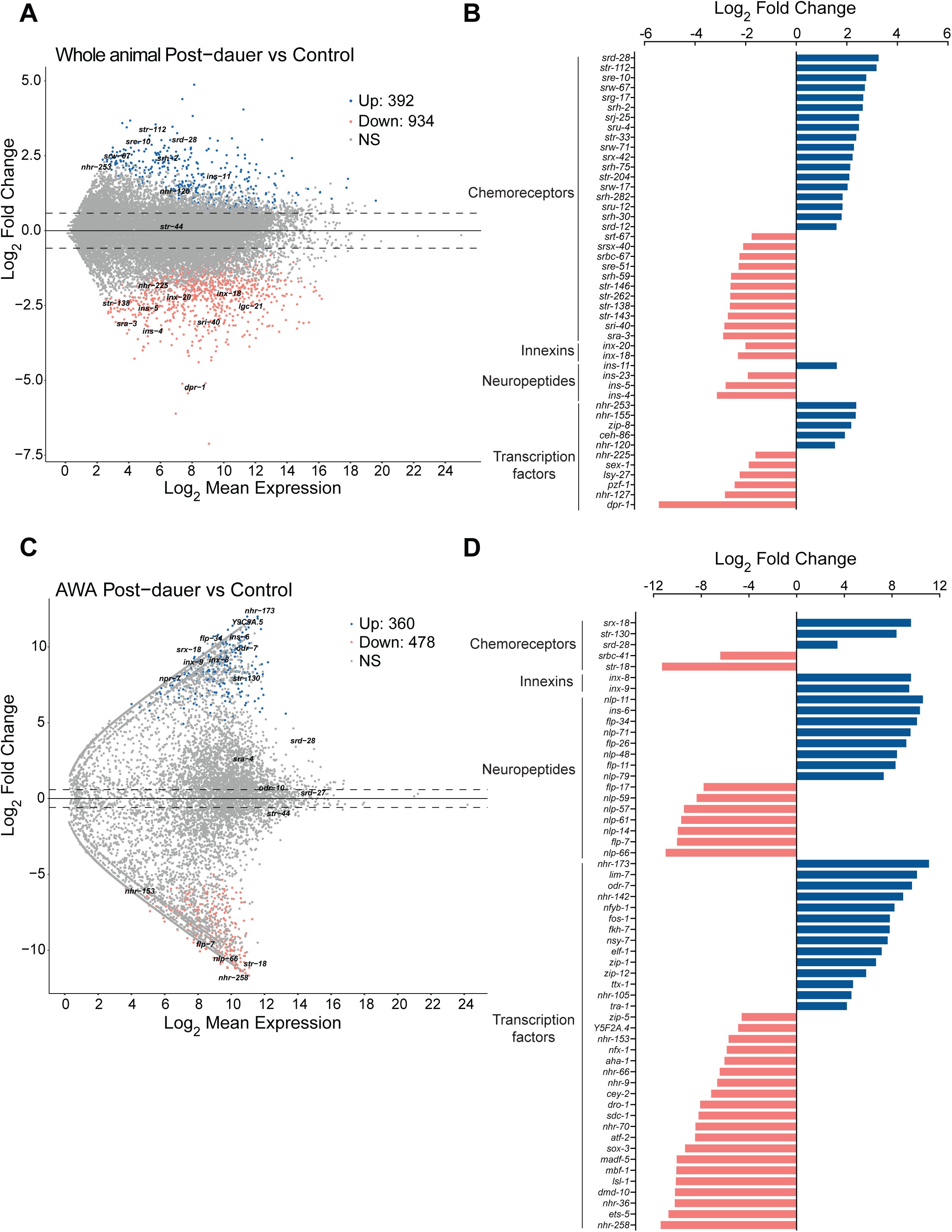
Gene expression is altered in AWA following passage through the dauer stage. **A)** MA plot showing enrichment and log2 fold changes of differentially expressed genes in PD vs control whole worm RNA-Seq libraries. The names of a subset of genes from different gene families are indicated. Up- and down-regulated genes were determined by differential expression analysis with a log2 fold change cut off > 1.5, padj < 0.05. **B)** Waterfall plot of differentially expressed genes from indicated gene families in control and PD whole animal RNA-Seq data. Up- and down-regulated genes were determined by differential expression analysis with a log2 fold change cut off > 1.5, padj < 0.05. **C)** MA plot showing enrichment and log2 fold changes of differentially expressed genes in PD vs control AWA RNA-Seq libraries. The names of a subset of genes from different gene families are indicated. Up- and down-regulated genes were determined by differential expression analysis with a log2 fold change cut off > 1.5, padj < 0.05. **D)** Waterfall plot of differentially expressed genes from indicated gene families in PD vs control AWA RNA-Seq libraries. Up- and down-regulated genes were determined by differential expression analysis with a log2 fold change cut off > 1.5, padj < 0.05.

In contrast to the gene expression changes in the whole animal dataset, the expression of only a small number of GPCRs appeared to be altered in PD vs control AWA neurons (Figure 5C-D, File S2). As expected, *odr-10* transcript levels were not significantly changed in PD adults. However, we noted that the expression of several neuropeptide genes and in particular, multiple transcription factors was significantly altered in PD vs control adults (Figure 5D, File S2). Affected transcription factors belong to multiple subfamilies including the greatly expanded nuclear hormone receptor family, members of which are predicted to be co-expressed in sensory neurons along with putative chemoreceptor GPCRs, and which have been suggested to act as receptors for external and internal cues (Sural and Hobert, 2021; Taylor et al., 2021). Together, these results indicate that passage through the dauer stage alters the expression of genes from multiple families, and that the expression of genes in individual sensory neurons such as AWA is also affected.

### The expression of multiple AWA-expressed chemoreceptor genes is upregulated in dauers and is maintained at higher levels in PD adults

As described above, while *odr-10* expression is not altered in PD adults, expression of this gene is transcriptionally upregulated in dauer larvae following which ciliary ODR-10::GFP protein levels are maintained at higher levels in PD adults. Since our RNA-Seq experiments indicated that the mRNA levels of only a small number of GPCR genes are altered in PD as compared to control AWA neurons, we tested whether additional AWA-expressed chemoreceptor genes are regulated similarly to *odr-10*.

Since we were unable to efficiently dissociate dauer larvae likely due to their modified cuticle (Cassada and Russell, 1975) and thus could not profile populations of sorted AWA neurons from dauer larvae, we instead examined the expression of a subset of endogenously tagged chemoreceptor genes. Similar to *odr-10*, mRNA levels of the putative AWA-expressed chemoreceptors *srd-27*, *srd-28*, and *str-44* were also not significantly upregulated in AWA in PD adults (Figure 5C-D, File S2) although *srd-28* was identified as a gene upregulated in PD adults in the whole animal dataset (Figure 5B). *srd-27, srd-28* and *str-44* endogenously tagged with *t2A::mNG* were specifically expressed bilaterally in the AWA olfactory neurons in L3 larvae and control adults (Figure 6A) (McLachlan et al., 2022). Expression of all three genes was upregulated in AWA in dauer larvae as compared to L3 larvae with *srd-28* and *str-44* showing stronger changes (Figure 6A-D). However, unlike *odr-10* whose expression remained restricted to AWA in all examined stages, all three chemoreceptor genes were expressed in additional neurons in the head in dauer larvae (Figure 6A). Expression in all neurons including in AWA was subsequently downregulated in PD adults relative to dauer larvae, although *srd-28*, *str-44*, and *srd-27* retained higher expression levels in PD as compared to control adults (Figure 6A-D). As in the case of *odr-10*, depletion of DAF-16 during dauer entry via auxin-induced degradation (Figure 4Ai) also decreased but did not abolish upregulation of *srd-28::t2A::mNG* expression in dauer larvae (Figure 6E). We conclude that the expression of multiple AWA-expressed chemoreceptor genes is likely transcriptionally upregulated in dauer larvae in part via DAF-16-dependent mechanisms, and that increased levels of a subset of these receptors may be maintained in PD adults possibly via non-transcriptional mechanisms.

**Figure 6.**
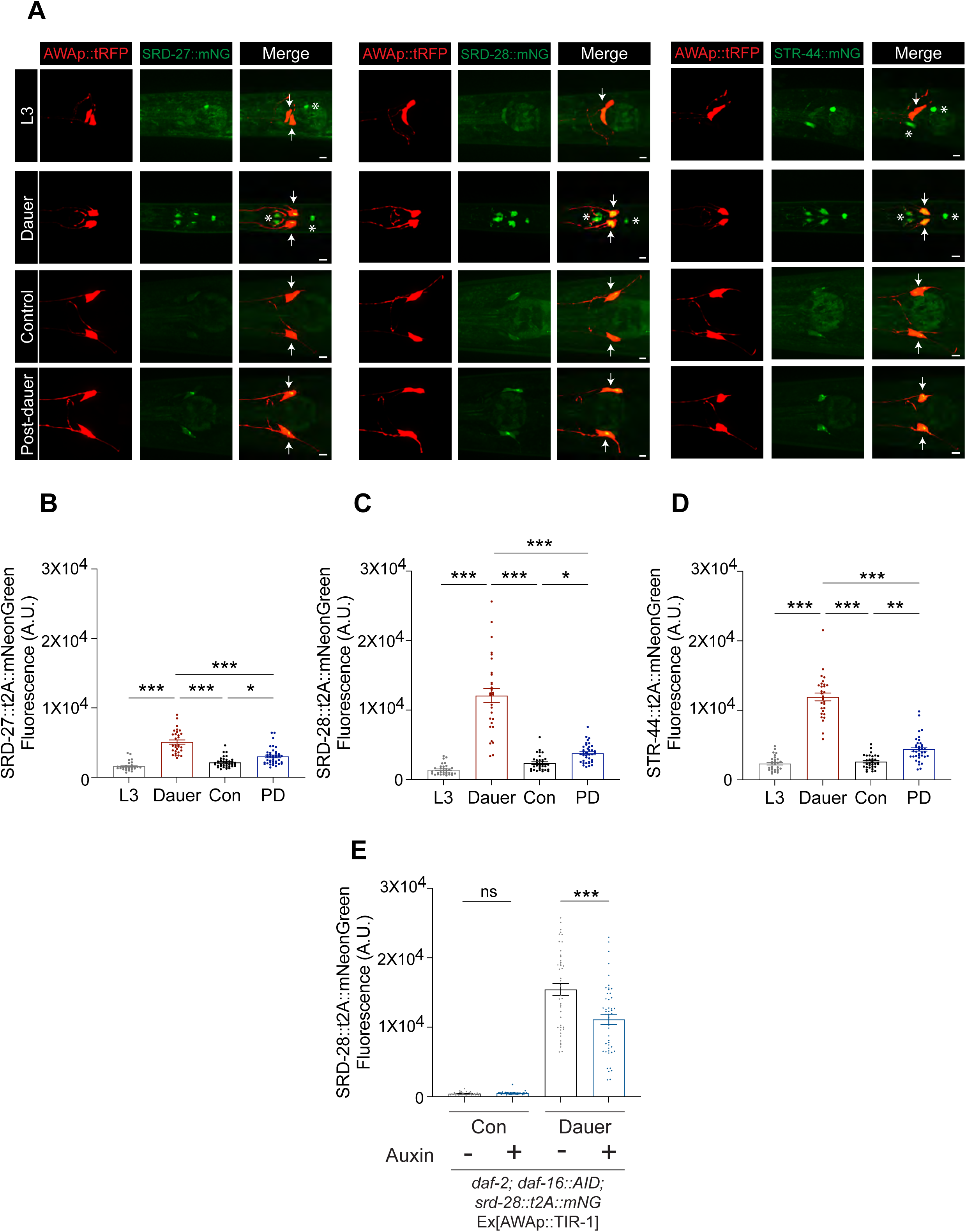
The expression of a subset of AWA-expressed chemoreceptor genes may be regulated similarly to *odr-10* upon dauer passage. **A)** Representative images of receptor::t2A::mNeonGreen expression from the corresponding endogenous loci in L3, dauer, control, and post-dauer adult stage animals. AWA neurons are marked via expression of *gpa-4Δ6*p::tagRFP. Arrows indicate AWA neurons; asterisks indicate ectopic expression in other cell types. Anterior is at left. Scale bars: 5 µm. **B,C,D)** Quantification of mean receptor::t2A::mNeonGreen fluorescence in AWA neurons in animals of at the indicated developmental stages. Each dot is a measurement from an individual neuron. Bars represent mean; error bars are SEM. n ≥ 28 neurons (≥ 18 animals). L3 and dauer larvae were imaged in parallel; 3 independent experiments. Control and PD adults were imaged in parallel; 3 independent experiments. *, **, *** indicates different at *P*<0.05, <0.01, and <0.001 (Kruskal-Wallis with Dunn’s multiple comparisons test); ns – not significant. **E)** Quantification of mean SRD-28::t2A::mNeonGreen fluorescence in AWA in *daf-2(e1368)* animals expressing *daf-16::mNeptune::AID* and *gpa-4Δ6*p*::TIR1* grown with or without 1 mM auxin. Animals were grown as indicated in the cartoon in Figure 4Ai. Each dot is a measurement from an individual neuron. Bars represent mean fluorescence, error bars are SEM. n ≥ 37 neurons (≥ 22 animals). Control and dauers grown on plates with and without 1 mM auxin plates were imaged in parallel; 3 independent experiments. *** indicates different at *P*<0.001 (two-tailed Welch’s t-test); ns – not significant.

## DISCUSSION

We show here that adult *C. elegans* hermaphrodites that transiently passed through the dauer developmental stage exhibit enhanced olfactory responses associated with food-seeking as compared to adult animals that bypassed this stage. Increased sensitivity to the odorant diacetyl is correlated with upregulated expression of the diacetyl receptor ODR-10 in dauer larvae, and higher levels of the ODR-10 protein in AWA olfactory cilia in PD adults. Via transcriptional profiling, we further show that levels of a subset of additional AWA-expressed chemoreceptors are also modulated by developmental stage and trajectory. Our results suggest that state- and experience-dependent cues are integrated to differentially modulate individual chemoreceptor levels in a single sensory neuron type likely via both transcriptional and post-transcriptional mechanisms, thereby providing a possible mechanism underlying developmental history-dependent olfactory behavioral plasticity.

Upon entry into the dauer stage, *odr-10* expression is transcriptionally upregulated in part via DAF-16. However, activation of DAF-16 alone appears to be insufficient to upregulate *odr-10* expression since *odr-10* expression is not upregulated in *daf-2* insulin receptor mutants that did not enter the dauer stage. Although we are unable to exclude the possibility that DAF-16 is only partially depleted in AWA upon auxin treatment, it is likely that additional factors including transcription factors such as the DAF-3 SMAD or the DAF-12 nuclear hormone receptor that act downstream of dauer-promoting hormonal signals also play a role in regulating *odr-10* expression (Aghayeva et al., 2021; Fielenbach and Antebi, 2008; Sims et al., 2016). Similarly, altered expression of innexins and other chemoreceptors in dauers were shown to be only partly DAF-16-dependent (Aghayeva et al., 2021). Upon exit from the dauer stage, *odr-10* expression is downregulated possibly due to inactivation of DAF-16 and the altered chromatin profile of these animals (Hall et al., 2010; Sims et al., 2016), but levels of ciliary ODR-10 protein continue to be maintained at higher levels than in control animals. Ciliary GPCRs are trafficked to the cilia base and further trafficked into and out of the cilium via diffusion and motor-driven transport (Mukhopadhyay et al., 2017; Nachury and Mick, 2019). We previously showed that ciliary trafficking of chemoreceptors in different *C. elegans* sensory neurons is regulated by multiple neuron- and receptor-specific mechanisms, and that these trafficking mechanisms are further modulated by sensory signaling (Brear et al., 2014; DiTirro et al., 2019). Sensory neuron cilia including those of AWA are extensively remodeled in dauers but their morphologies appear to be restored to those resembling those in control animals in PD adults (Albert and Riddle, 1983; Britz et al., 2021) (this work). We propose that one or more ciliary trafficking mechanisms are altered in PD animals resulting in increased ODR-10 protein trafficking into, or decreased removal from, the AWA cilia in PD animals. In addition to transcriptional changes in chemoreceptor expression, regulated trafficking of ciliary GPCRs provides an additional mechanism to fine tune sensory responses as a consequence of developmental experience.

Why do PD adults enhance their responses to food-related odors? Stress, including starvation, experienced during early larval stages may indicate that food availability is unreliable. Increased attraction to food in PD adults may be a bet-hedging strategy that optimizes growth and survival in a variable environment. However, although animals also retain a cellular memory of starvation-induced L1 arrest (Jobson et al., 2015; Webster et al., 2018), this arrest does not appear to alter examined olfactory behaviors in adults, suggesting that stress assessed during a distinct period in development and/or dauer entry is required to drive behavioral plasticity in ensuing adults. Interestingly, we and others previously showed that expression of the *osm-9* TRPV channel gene is strongly downregulated in the ADL nociceptive but not AWA neurons in PD adults, resulting in decreased responses to the ADL-sensed aversive ascr#3 pheromone (Sims et al., 2016). Decreased aversion to this pheromone by PD adults has been suggested to be a mechanism that inhibits dispersion in a crowded environment and promotes outcrossing (Sims et al., 2016). Coordinated differential modulation of responses in defined subsets of chemosensory neurons likely allow PD adults to optimize survival and reproduction.

In contrast to the main olfactory systems of vertebrates in which each olfactory sensory neuron expresses one or very few olfactory receptors (Chess et al., 1994; Ressler et al., 1993; Vassar et al., 1993), the co-expression of as many as 100 chemoreceptors in each chemosensory neuron type in *C. elegans* (Taylor et al., 2021; Troemel et al., 1995; Vidal et al., 2018) raises a unique challenge for this organism. Since co-expressed chemoreceptors are likely tuned to distinct odorants, modulating synaptic transmission from a single chemosensory neuron in *C. elegans* as a function of internal state would be expected to coordinately alter responses to a broad range of chemicals sensed by that neuron type. Regulation of individual chemoreceptors instead enables the animal to precisely target and modulate defined chemosensory behaviors. Consistent with this notion, expression of chemoreceptor genes is subject to complex modes of regulation. *odr-10* expression is higher in hermaphrodites and juvenile male larvae than in adult males, and expression is upregulated in adults of both sexes upon starvation (Ryan et al., 2014; Wexler et al., 2020). The expression of multiple chemoreceptors in many sensory neuron types is dramatically altered in dauer larvae and subsequently further modulated in PD animals, whereas the expression of other receptors is sexually dimorphic (Nolan et al., 2002; Peckol et al., 1999; Troemel et al., 1995; Vidal et al., 2018). In the ADL nociceptive neurons, expression of the *srh-234* chemoreceptor has been shown to be regulated via cell-autonomous and non cell-autonomous mechanisms (Gruner et al., 2016; Gruner et al., 2014). Thus, the expression of each chemoreceptor gene may be modulated by multiple regulatory modules that act combinatorially to integrate distinct inputs and appropriately calibrate the behavioral response.

Individual chemosensory neurons in *Aedes aegypti* also express multiple receptors from different receptor subfamilies (Younger et al., 2020). The expression of subsets of these receptors has been shown to be modulated by feeding and reproductive state although whether receptor expression is causative to altered olfactory behavioral profiles is unclear (Matthews et al., 2016). In rodents, a subset of specialized olfactory neurons in rodents express multiple receptors of the MS4A family, each of which responds to ethologically relevant chemical stimuli (Greer et al., 2016). It will be interesting to assess whether expression of these receptors is also subject to extensive state-dependent modulation, and whether this modulation drives sensory behavioral plasticity as a function of current and past external and internal conditions.

## DATA AVAILABILITY STATEMENT

All plasmids used in this work are listed in Table S1. All strains used in this work are listed in Table S2. Strains and plasmids are available upon request. Data underlying this article are available at https://github.com/SenguptaLab/PDplasticity. Sequencing files of the RNA-Seq experiments have been deposited in the Array Express database at EMBL-EBI (www.ebi.ac.uk/arrayexpress) under accession number E-MTAB-11823.

## ACKNOWLEDGEMENTS

We thank Oliver Hobert and Mario de Bono for reagents and experimental advice, David Miller, Nicolas Henderson and Dylan Ma for assistance with cell sorting, Albert Yu and Ines Patop for assistance with analyses of RNA-Seq data, Bradly Stone for assistance with statistical analyses, and the *Caenorhabditis* Genetics Center for strains. We are grateful to members of the Sengupta lab for advice and members of the Sengupta lab, Doug Portman and Sarah Hall for critical comments on the manuscript.

## SUPPLEMENTAL FIGURE LEGENDS

**Figure S1.**
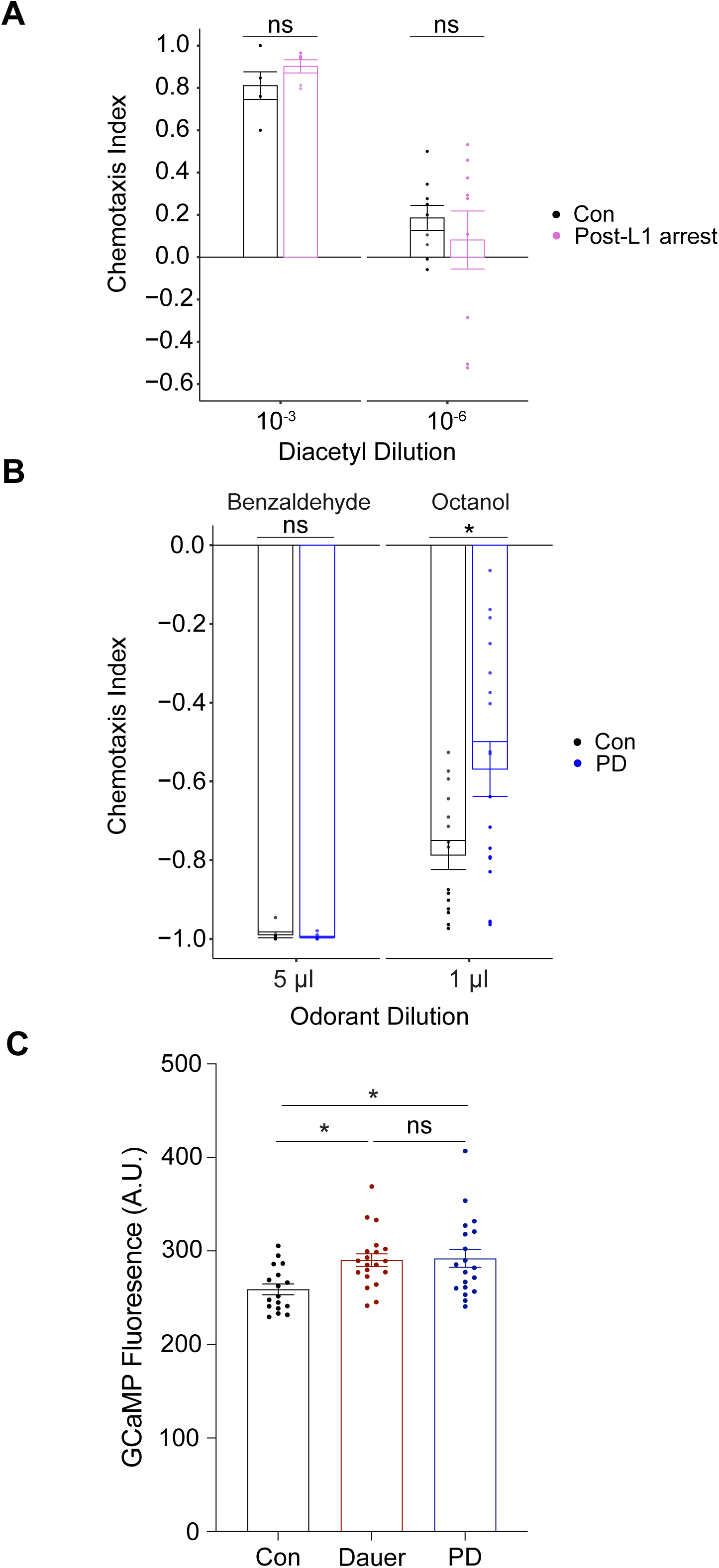
Adults that experienced L1 arrest do not exhibit increased diacetyl responses. **A)** Behavioral responses of wild-type control (con) and post-L1 (post-L1 arrest) arrested adults to indicated dilutions of diacetyl. Each dot is the chemotaxis index of a single assay plate containing ∼50-300 adult hermaphrodites. Bars represent the mean; error bars are SEM. The behaviors of control and post-L1 arrested animals were assayed in parallel in duplicate; ≥3 independent experiments; ns – not significant (two-tailed Welch’s t-test). **B)** Behavioral responses of wild-type control and PD adults to the indicated concentrations of aversive volatile odorants. Each dot represents the chemotaxis index of a single assay plate containing ∼50-300 adult hermaphrodites. Bars represent the mean; error bars are SEM. Control and PD behaviors were assessed in parallel; ≥ 3 independent experiments. * indicates different at *P*<0.05 (two-tailed Welch’s t-test); ns – not significant. **C)** Baseline GCaMP2.2b fluorescence in AWA soma in the indicated wild-type animals. Baseline measurements were collected from experiments reported in Figures 1D and 3F. Each dot is the measurement from a single neuron. Bars represent the mean; error bars are SEM. n ≥ 17 animals (1 neuron per animal). * indicates different at *P*<0.05 (one-way ANOVA with Tukey’s multiple comparisons test); ns – not significant.

**Figure S2.**
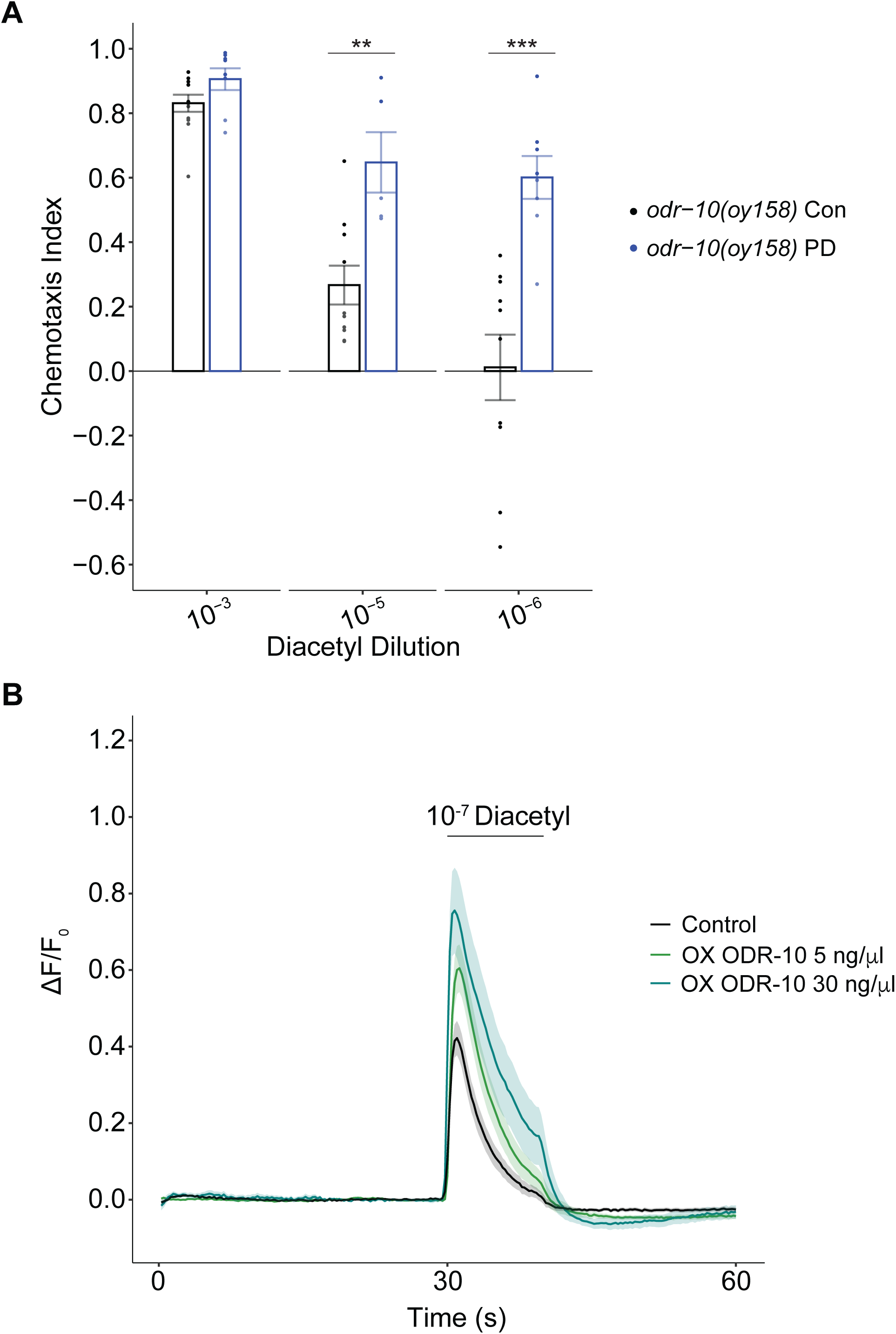
An endogenously tagged *odr-10::gfp_11_* strain retains dauer passage-dependent olfactory behavioral plasticity as adults. **A)** Behavioral responses of *odr-10(oy158)* control and PD animals expressing *odr-10::gfp_11_* from the endogenous *odr-10* locus to the indicated dilutions of diacetyl. Each dot represents the chemotaxis index of a single assay plate containing ∼50-300 adult hermaphrodites. Bars represent the mean; error bars are SEM. Chemotaxis assays at each concentration were performed in parallel over at least three days. ** and *** indicate different at each concentration at *P*<0.01 and 0.001, respectively (two-tailed t-test with Welch’s correction). **B)** Average changes in GCaMP2.2b fluorescence in AWA soma in control animals of the indicated genotypes to a 10 sec pulse of 10^-7^ diacetyl. Shaded regions indicate SEM. n ≥ 16 animals (1 neuron per animal) each.

**Figure S3.**
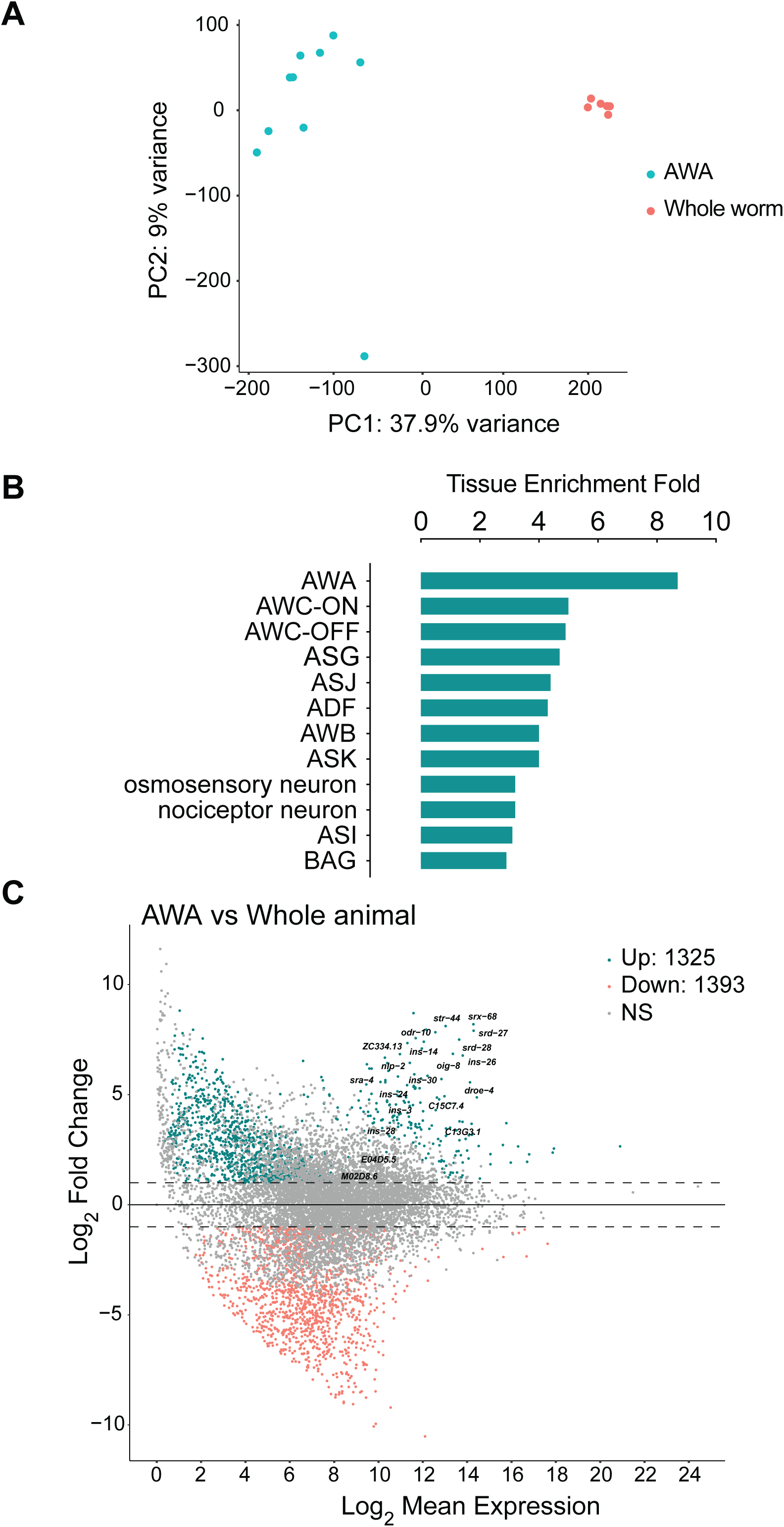
Transcriptional profiling of sorted populations of AWA neurons. **A)** PCA clustering of RNA-Seq libraries from sorted populations of AWA neurons and dissociated cells from whole animals based on the 10,000 most differentially expressed genes. **B)** Tissue Enrichment Analysis with the web-based Tissue Enrichment Analysis tool (https://www.wormbase.org/tools/enrichment/tea/tea.cgi) (Angeles-Albores et al., 2016) shows enrichment of AWA-expressed genes in AWA RNA-Seq libraries. log2 fold change cut off > 2, padj < 0.05. **C)** MA plot showing differentially expressed genes in RNA-Seq data from FACS-sorted populations of control and PD AWA neurons as compared to dissociated but unsorted cells from control and PD whole animals. 20 of the most highly AWA-expressed genes (Taylor et al., 2021) are indicated. Up- and down-regulated genes were determined by differential expression analysis with a log2 fold change cut off > 2, padj < 0.05.

**File S1.** RNA-Seq data of differentially expressed genes from control and PD whole animals.

**File S2.** RNA-Seq data of differentially expressed genes from sorted control and PD AWA neurons.

**Table S1.** Plasmids used in this work.

| Plasmid | Construct | Source | Relevant Figures |
| --- | --- | --- | --- |
| PSAB1144 | *gpa-4Δ6p::mks-5::rfp* | This work | 2C, 2D, S2A, 3C, 3D, 4D |
| PSAB1280 | *gpa-4Δ6p::gfp<sub>1-10</sub>* | This work | 2C, 2D, S2A, 3C, 3D, 4D |
| PSAB1281 | *gpa-4Δ6p::odr-10::tagRFP* | This work | 2E, 2F, S2B |
| PSAB1282 | *gpa-4Δ6p::tagRFP* | This work | 6A, 6B, 6C, 6D |
| PSAB1283 | *gpa-4Δ6p::tir1::SL2::mScarlet* | This work | 4A, 4B, 6E |

**Table S2.**
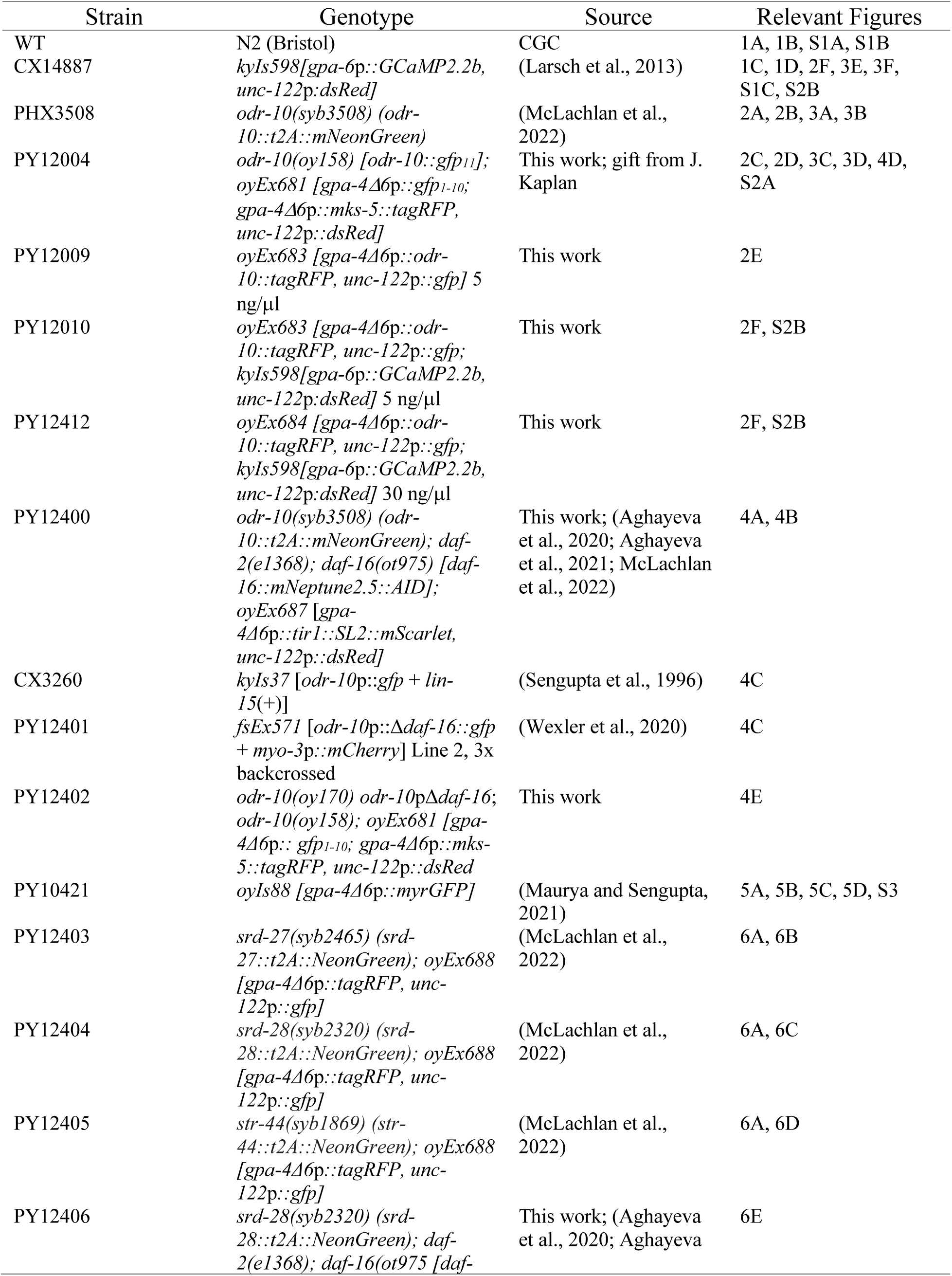
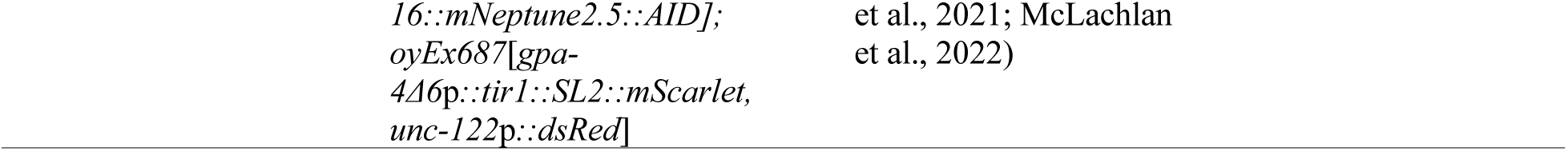
Strains used in this work.

